# DNA-stabilized silver nanoclusters as specific, ratiometric fluorescent dopamine sensors

**DOI:** 10.1101/205591

**Authors:** Jackson T. Del Bonis-O’Donnell, Ami Thakrar, Jeremy Wain Hirschberg, Daniel Vong, Bridget N. Queenan, Deborah K. Fygenson, Sumita Pennathur

**Affiliations:** Department of Mechanical Engineering, University of California, Santa Barbara, CA, USA; Neuroscience Research Institute, University of California, Santa Barbara, CA, USA; Department of Physics and Program in Biomolecular Engineering, University of California, Santa Barbara, CA, USA

**Keywords:** Neurotransmitter, dopamine, biosensor, DNA-stabilized silver nanoclusters

## Abstract

Neurotransmitters are small molecules that orchestrate complex patterns of brain activity. Unfortunately, there exist few sensors capable of directly detecting individual neurotransmitters. Those sensors that do exist are either unspecific or fail to capture the temporal or spatial dynamics of neurotransmitter release. DNA-stabilized silver nanoclusters (DNA-AgNCs) are a new class of biocompatible, fluorescent nanostructures that have recently been demonstrated to offer promise as biosensors. In this work, we identify two different DNA sequences which form dopamine-sensitive nanoclusters. We demonstrate that each sequence supports two distinct DNA-AgNCs capable of providing specific, ratiometric fluorescent sensing of dopamine concentration *in vitro*. DNA-Ag nanoclusters therefore offer a novel, low-cost approach to quantification of dopamine, creating the potential for real-time monitoring *in vivo*.

## INTRODUCTION

Neurotransmitters (NTs) are small molecules used in signaling pathways throughout the nervous system. The dynamic patterns of neurotransmitter signaling are critical to brain function and tightly regulated, both temporally and spatially. Improper release, reuptake, and/or degradation of neurotransmitters has been implicated in a wide range of neurodevelopmental, neurological and neurodegenerative disorders, from epilepsy^1^ to depression^2^ to Parkinson’s disease.^3^ Monitoring patterns of neurotransmitter release in the brain would provide fundamental insights into brain function and could serve as a diagnostic tool for maladaptive mental states and diseases. However, the ability to quantify specific endogenous neurotransmitters is limited.

Current methods to detect neurotransmitters are predominantly electrochemical. Certain molecules, including the biogenic amines (which include dopamine, epinephrine, serotonin, and ascorbic acid) can be oxidized by an applied potential, allowing nearby microelectrodes to detect the chemical reaction as an electrical signal.^4^ The time resolution of these methods is as fast as the electrochemical reaction, but the specificity is low: the applied potential will oxidize any biogenic amine present. Constant-potential amperometry has been adapted to increase performance; the resulting fast-scan cyclic voltammetry is now coupled with microdialysis to detect the biogenic amines^5–7^ (as well as oxygen and pH^8^) more reliably *in vivo*.

However, while electrochemical methods provide the millimeter spatial and millisecond temporal resolution adequate for neuro-signal sensing,^9^ their lack of specificity severely limits signal interpretation. Furthermore, the family of electrochemical techniques is simply not applicable to other small molecules, including the major neurotransmitters, glutamate and GABA. Further research is needed to enable specific and reliable detection of individual neurotransmitters with appropriate spatiotemporal resolution.

The use of optically active nanomaterials for specific binding and fluorescent detection is an emerging and promising approach to *in vivo* sensing of biomolecules.^10–12^ However, widespread utility will be limited if complex design or costly synthesis is required for each biological target. DNA-stabilized fluorescent silver nanoclusters (DNA-AgNCs) are a new class of fluorescent nanostructure formed when a particular sequence of single-stranded DNA stabilizes a few-atom cluster of silver.^13,14^ DNA-AgNCs offer certain key advantages as biosensors. DNA-AgNCs are relatively cheap and easy to synthesize, and are biocompatible.^15^ The fluorescence of a DNA-AgNC is largely determined by the stabilizing DNA sequence and therefore readily manipulated^16–18^. Perturbations to the local environment or DNA conformation of DNA-AgNCs can produce measureable changes in their emissive properties.^16,19^ These combined properties have led to the use of DNA-AgNC probes as novel fluorescent sensors for metal ions and DNA sequences in complex biological matrices.^20–22^ We theorized that DNA-stabilized AgNC probes could serve as fluorescent sensors for biologically relevant small molecules, such as neurotransmitters.

Here we demonstrate that DNA-AgNCs can be specifically sensitive for dopamine. We identify two distinct DNA-AgNCs which show preference for dopamine over other neurotransmitters. These two DNA probes enable two classes of dopamine sensing: one provides a “turn-on” fluorescent sensor in the presence of dopamine and the other provides a “turn-off” fluorescent sensor. Importantly, we determined that these two DNA sequences each produced silver nanocluster populations with distinct spectral populations. The nanocluster populations responded differentially to dopamine, providing a ratiometric measure of dopamine concentration *in vitro*. We validated that these DNA-AgNC probes were responsive to dopamine, but insensitive to other common neurotransmitters, related catecholamines, and known competitors. We therefore present DNA-AgNCs as novel, biocompatible sensors of dopamine with potential for quantitative, ratiometric, optical *in vivo* sensing of neurotransmitters.

## RESULTS & DISCUSSION

We hypothesized that DNA-stabilized silver nanoclusters could be used as sensors of small molecule neurotransmitters. We selected six DNA ‘template’ sequences (sequences S1-S6, ***Table 1***) known to stabilize silver atoms into fluorescent nanoclusters, ^16,17,23,24^ and synthesized AgNCs on each using a standard approach: addition of AgNO_3_ followed by reduction using NaBH_4_. We first determined the basal fluorescence of the DNA-AgNC created by each probe (***Figure 1a***). In these initial screens, we used UV excitation (260 nm) because it universally excites fluorescent DNA-AgNCs regardless of their specific visible excitation spectra.^25^ In the absence of neurotransmitter, all six template sequences produced bright, stable DNA-AgNCs with emission peaks between 500-800 nm (***Figure 1a***) consistent with previous reports (see ***Figure S1*** and ***Supporting Information*** for spectral details).

**Figure 1.**
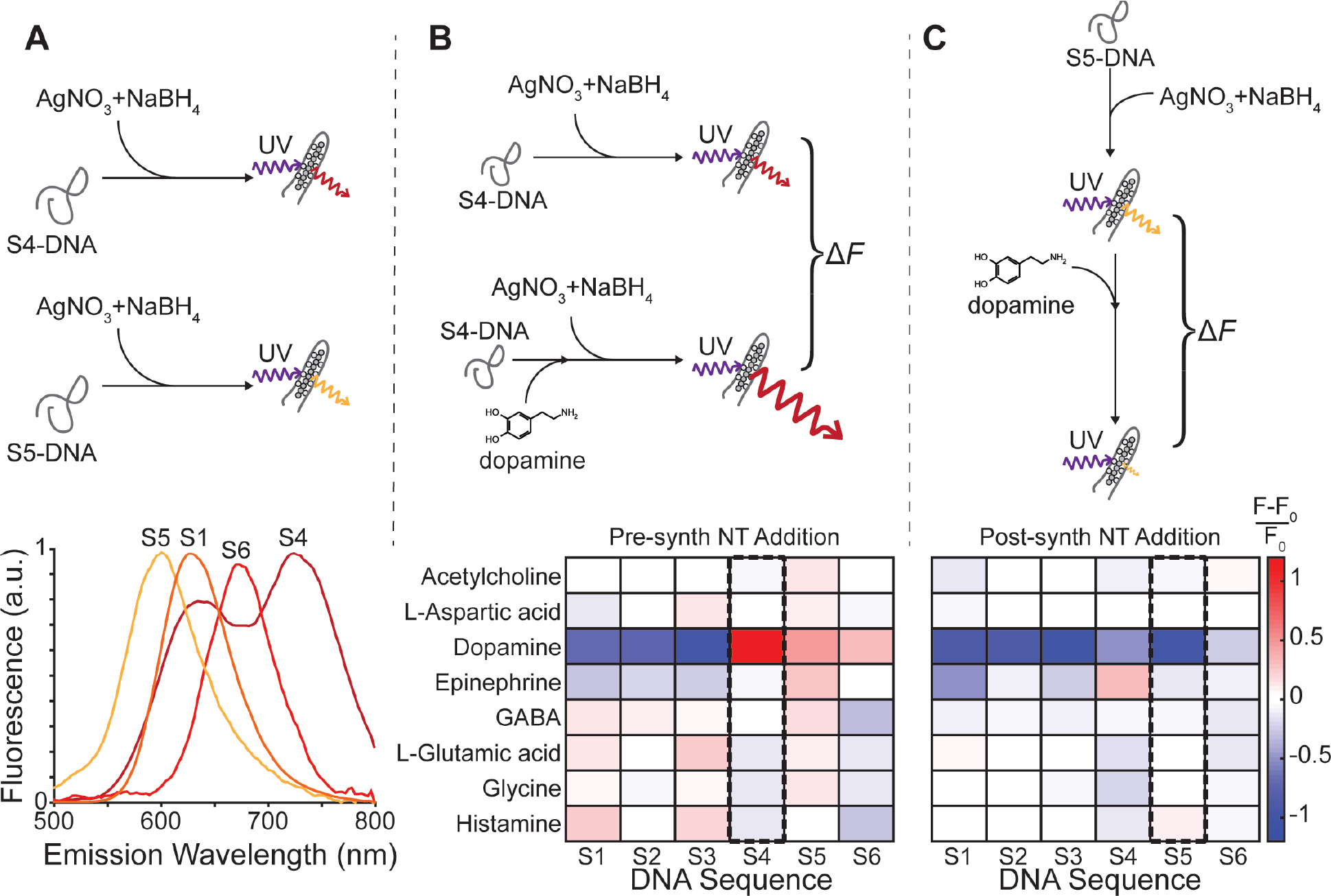
Changes in DNA-AgNC fluorescence in the presence of neurotransmitter. *Top panels:* Schematic representation of experimental approach. **(a)** We synthesized silver nanoclusters (AgNCs) using 6 DNA sequences (S1-S6) and determined the fluorescence emission spectra of each nanocluster. For these initial screens, we used UV excitation (260 nm, purple) because it universally excites fluorescent DNA-AgNCs regardless of their specific visible excitation spectra.^25^ *Bottom:* Fluorescence emission spectra of each DNA-AgNC. DNA sequences S2 and S3 had emission spectra similar to those of S5 and were thus omitted for clarity. **(b-c)** Peak fluorescence emission response of the 6 DNA-AgNCs (x-axis, [DNA] = 12.5 μM) in the presence of 8 neurotransmitters (NTs, y-axis, [NT] = 50 μM). Baseline fluorescence for each sequence (white) was established by exciting each DNA-AgNC in the absence of NT. Sensitivity of the nanoclusters to NT was assayed under two conditions: neurotransmitters were added either before **(b)** or after **(c)** the assembly of fluorescent nanoclusters. Relative fluorescence was then quantified for each sequence: red indicates increased while blue indicates decreased fluorescence in the presence of NT relative to baseline DNA-AgNC fluorescence for each DNA sequence. Uniquely, DNA S4 nanocluster fluorescence dramatically increased when dopamine was present during nanocluster synthesis (boxed column S4 and schematic in **b**). Conversely, DNA S5 nanocluster fluorescence dramatically decreased when dopamine was added after synthesis (boxed column S5 and schematic in **c**).

**Table 1.**
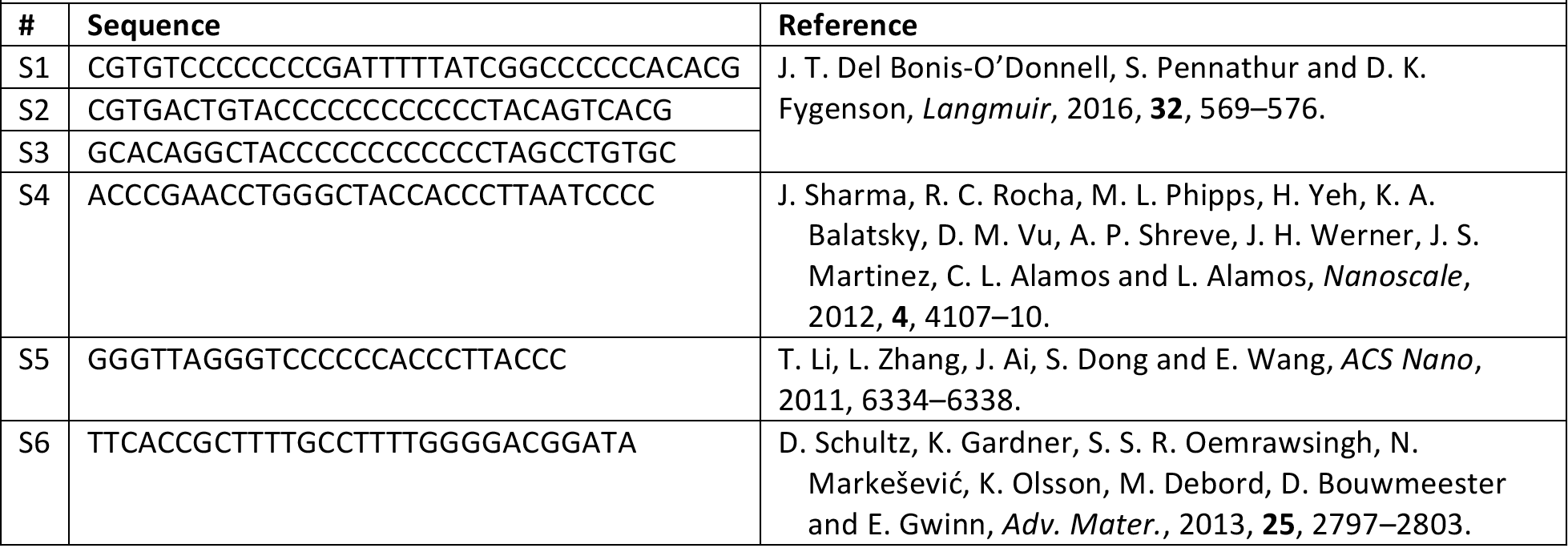
Oligonucleotides sequences.

To identify potential DNA-AgNC sensors for neurotransmitter (NT) detection, we screened a library of eight well-studied NTs (acetylcholine, L-aspartic acid, dopamine, epinephrine, GABA, L-glutamic acid, glycine, and histamine, see ***Supplementary Figure S5*** for chemical structures) against the six DNA-AgNCs (***Figure 1b-c***). We used the peak emission of each probe (represented as white) to gauge the change in intensity elicited by the presence of NT either during or after nanocluster synthesis. We first tested whether emission intensity was altered by the presence of NT during synthesis (***Figure 1b***). Of all the NTs tested, dopamine produced the greatest changes in DNA-AgNC fluorescence. When nanoclusters were assembled in the presence of dopamine, samples containing templates S1-3 fluoresced less, while templates S4-6 fluoresced more than DNA-AgNCs assembled in the absence of neurotransmitter. We then determined whether the fluorescence of assembled DNA-AgNC nanoclusters was altered by the addition of NTs after synthesis (***Figure 1c***). Most DNA-AgNCs showed mild reductions in fluorescence when NT was added. However, when dopamine was added, the fluorescence of assembled DNA-AgNCs formed using sequences S1, S2, S3 and S5 decreased profoundly.

Of the six DNA-AgNCs tested, two showed promise for sensor development. The fluorescence of nanoclusters assembled using DNA sequence S4 increased dramatically in the presence of dopamine during synthesis, though it was largely unaffected by other NTs (**Figure 1b**, boxed column S4). Conversely, the fluorescence of assembled DNA S5 nanoclusters declined dramatically upon addition of dopamine, but was unchanged by other NTs (**Figure 1c**, boxed column S5). The fluorescence emission of DNA S2-AgNCs and DNA S3-AgNCs were similar to DNA S5-AgNCs, but DNA S5-AgNCs exhibited a stronger quenching response and was more selective to dopamine. We therefore focused our efforts on the two strongly dopamine-responsive DNA sequences, S4 and S5.

### DNA S4 silver nanoclusters as ratiometric, “turn-on” dopamine sensors

Our initial screens identified neurotransmitter-responsive DNA-AgNCs using universal UV excitation at 260 nm and monitoring emission at the specific wavelength in the visible spectrum where intensity was greatest in the absence of NT. For our focused studies, we tailored the excitation wavelength to the individual construct and collected emission spectra. We first turned our attention to the silver nanoclusters assembled by DNA sequence S4. Collecting a full emission spectrum from the DNA S4-AgNCs using 525 nm excitation revealed an emission peak at 625 nm that more than doubled in intensity when dopamine was present (***Figure 2c***) but was barely changed by the other compounds tested (***Figure 2b***).

**Figure 2.**
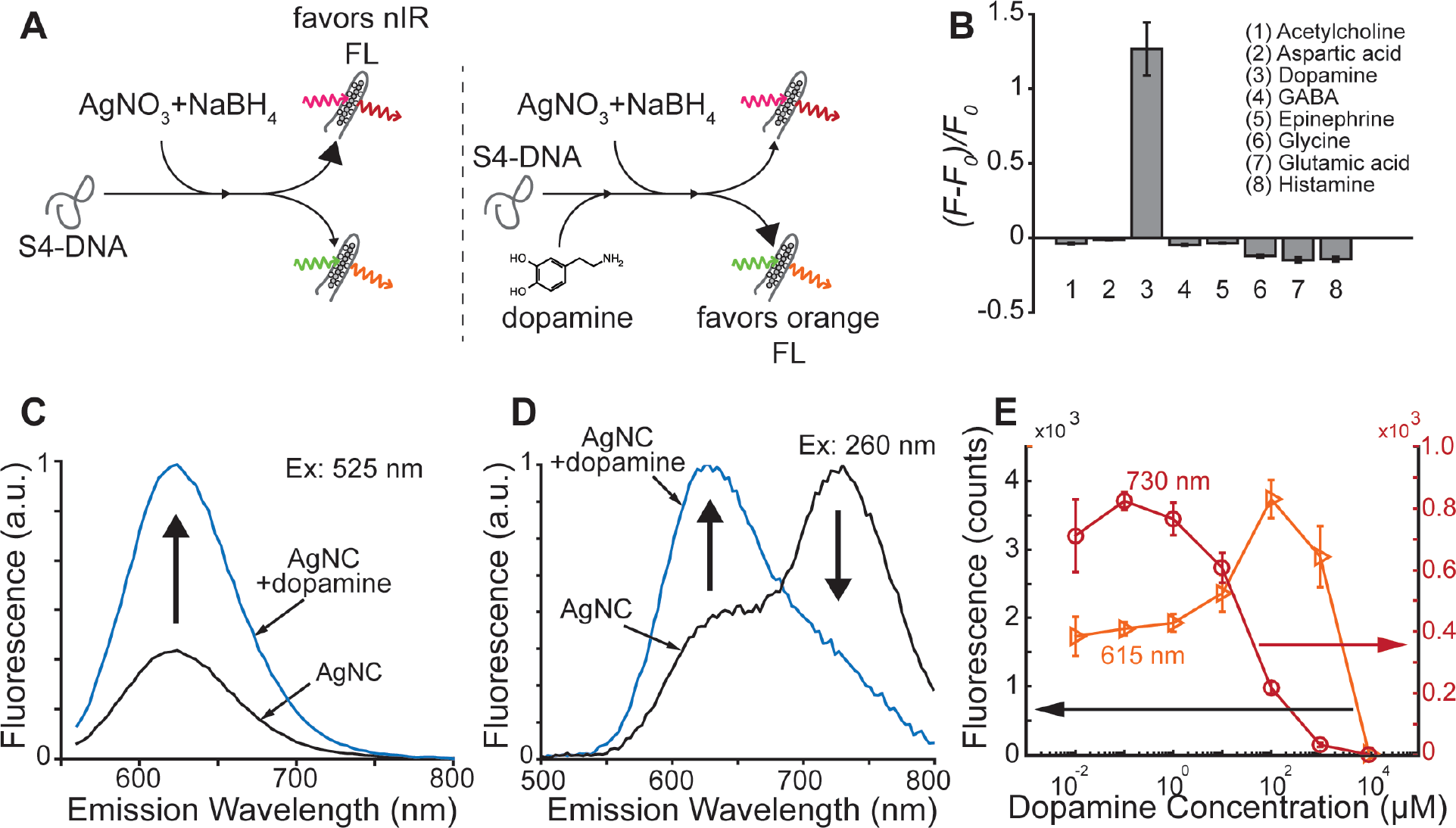
DNA S4 silver nanoclusters acts as ratiometric, “turn-on” dopamine sensors. **(a)** Schematic summary of experimental results. DNA sequence S4 produces two spectrally distinct AgNC populations. Formation of the long-wavelength (nIR) population (640 nm excitation, 730 nm emission, top) is favored in the absence of dopamine, while formation of the short-wavelength (visible) population (535 nm excitation, 615 nm emission, bottom) is favored in the presence of dopamine. **(b)** Change in fluorescence emission intensity (Δƒ = [F-F_0_]/F_0_) of AgNCs synthesized using DNA S4 in the presence of eight different NTs ([DNA] = 12.5μM, [NT] = 50 μM). Fluorescence emission was collected at 610 nm using 525 nm excitation (peak excitation for the S4-AgNCs). Error bars indicate the standard deviation of n=3 replicate experiments. **(c-d)** Fluorescence emission spectra of S4-AgNCs assembled in the absence (gray) and presence (blue) of dopamine under 525 nm **(c)** and 260 nm **(d)** excitation. In the presence of dopamine, intensity at the lower wavelength emission peak (615 nm) more than doubles while intensity at the higher wavelength peak (730 nm emission) drops a comparable amount. **(e)** Fluorescence emission response of the two spectrally distinct AgNCs generated by S4 when different concentrations of dopamine are added prior to cluster synthesis. Orange triangles: 535 nm excitation, 615 nm emission nanoclusters; red circles: 640 nm excitation, 730 nm emission nanoclusters. One of the two AgNCs species (730 nm emission, red) was significantly quenched by moderate amounts of dopamine, while the other species (615 nm emission, orange) was enhanced by moderate amounts of dopamine.

The S4 template was also exceptional in that its excitation with UV (260 nm) illumination clearly revealed two spectrally distinct clusters (***Figure 2d***). When illuminated with visible light at their respective peak excitation wavelengths, one cluster has a peak emission wavelength of 625 nm (as reported previously^23^) and the other has peak emission at 730 nm (not previously reported) (***Figure 1a, Figure 2d***). The presence of dopamine during nanocluster formation dramatically increases the 625 nm peak, while decreasing the 730 nm peak (***Figure 2d***).

We systematically tested the effect of dopamine concentration on the formation of these two spectrally distinct populations. When DNA-AgNC were templated with sequence S4 in the absence of dopamine, the formation of the near infrared-emitting (730 nm) population was favored (***Figure 2d-e***). However, as dopamine concentration increased from 1 μM to 100 μM, nIR fluorescence (730 nm emission) decreased approximately 66% and was effectively quenched at 1 mM dopamine (***Figure 2e***). In contrast, red fluorescence (collected at 615 nm) doubled in intensity as dopamine concentration increased from 1 μM to 100 μM (***Figure 2e***) and was not quenched until 10 mM dopamine. It should be noted that the fluorescence of both constructs decreased above 100 μM dopamine and, at these high concentrations, we observed silver nanoparticle formation and aggregation, which may have been induced directly by dopamine,^12^ or by competition for silver between the bases and the excess chloride present from using dopamine-HCl salt in our preparation.

Although nanoparticle formation and aggregation prevent quantitative interpretation at dopamine concentrations over 100 μM (***Figure 2e***, ***Figure S3***), at concentrations below 100 μM, the two emission peaks together provide a reliable, ratiometric signal (F_red_/F_nIR_) that changes by nearly two orders of magnitude (***Figure S3***), a greater dynamic range than either signal alone and comparable to the range of dopamine concentrations it reflects. We therefore propose that the relative fluorescence of the two spectrally distinct and differentially dopamine-sensitive DNA S4-AgNC populations (***Figure 2a***) can provide a robust, quantitative and sensitive measure of dopamine concentration *in vitro*.

### DNA S5 silver nanoclusters as ratiometric, “turn-off” dopamine sensors

We next turned our attention to nanoclusters stabilized by DNA sequence S5. In our initial screens, we observed that, when excited with UV light (260 nm), S5-AgNCs had a peak emission wavelength of 600 nm (***Figure 1a***) and a fluorescence intensity that was largely unaffected by the addition of most neurotransmitters (***Figure 1c*,** S5 column). However, when dopamine was added, DNA S5-AgNCs exhibited a ~80% decrease in fluorescence intensity (***Figure 1c***). Based on these observations, we scanned excitation wavelengths while holding emission fixed at 600 nm and identified 535 nm as the peak visible excitation wavelength. We then re-assessed the DNA S5-AgNC response to neurotransmitters. Encouragingly, addition of dopamine reduced fluorescence at the 600 nm emission peak even more under 535 nm excitation than under 260 nm excitation, diminishing by nearly two orders of magnitude (96% decrease, ***Figure 3b-c***). Meanwhile, changes in fluorescence in response to the other NTs were less than 11% (***Figure 3b***).

**Figure 3.**
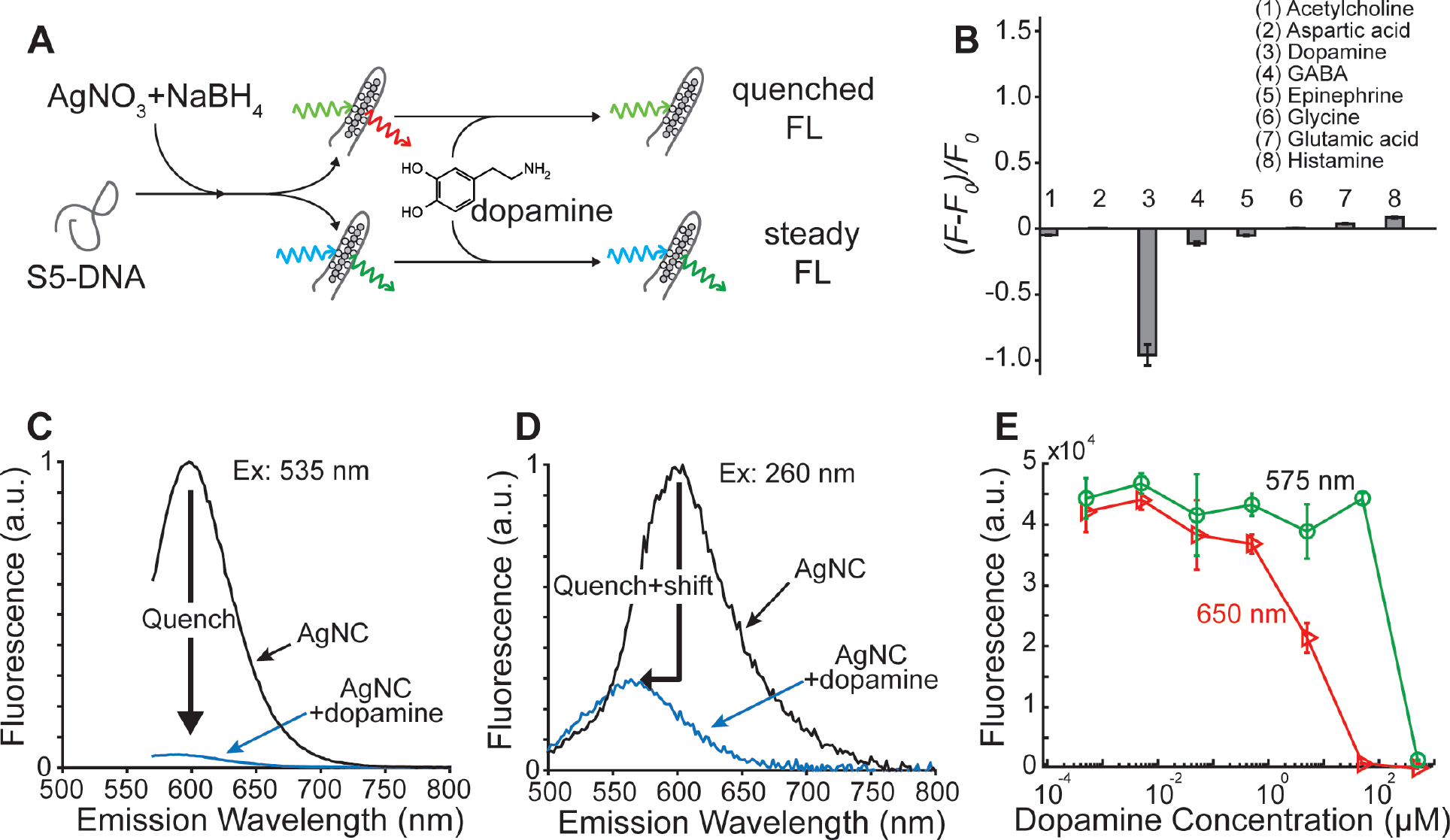
DNA S5 silver nanoclusters acts as ratiometric, “turn-off” dopamine sensors. **(a)** Schematic summary of experimental results. DNA sequence S5 produces two spectrally distinct AgNC populations. Fluorescence of the long-wavelength population (580 nm excitation, 650 nm emission, top) is quenched by dopamine in the 1-10 μ.M range, while that of the short-wavelength population (495 nm excitation, 575 nm emission, bottom) is not significantly affected by dopamine concentrations up to 50 μM. **(b)** Change in fluorescence emission intensity (Δƒ = [F-F_0_]/F_0_) of assembled DNA S5-stabilized AgNCs upon addition of eight different NTs ([DNA] = 12.5 μM, [NT] = 50 μM). Fluorescence emission was collected at 600 nm using 535 nm excitation (peak visible excitation for the S5-AgNCs). Error bars indicate the standard deviation of n=3 replicate experiments. **(c-d)** Fluorescence emission spectra of assembled S5-AgNCs without (black) and with dopamine (blue) under 535 nm **(c)** and 260 nm **(d)** illumination. Note the decrease in overall intensity and the shift in peak emission to shorter wavelengths under UV illumination **(d)**. **(e)** Fluorescence emission response of the two spectrally distinct AgNCs generated by S5 for different concentrations of dopamine added after cluster synthesis. Green circles: 495 nm excitation and 575 nm emission; red triangles, 580 nm excitation and 650 nm emission. Only one of the two AgNCs species (650 nm emission, red) was significantly quenched by moderate amounts of dopamine. [DNA]=1.25 μM.

Upon closer inspection, we noticed this response to dopamine was actually a compound effect: under 260 nm excitation adding dopamine to S5-AgNCs both decreased emission at 600 nm and shifted the peak emission from 600 nm to 560 nm (***Figure 3d***). Full excitation and emission spectra of the DNA S5-AgNCs revealed two cluster populations with distinct spectral properties: one with a peak emission at 650 nm (580 nm excitation) and another with a peak emission at 575 nm (495 nm excitation) (***Figure S4***). These populations were not resolved using 260 nm excitation (***Figure 1a, Figure 3d***) and their sum produced an apparent emission peak at 600 nm.

We examined the dopamine sensitivity of these two spectrally distinct nanocluster populations generated by DNA sequence S5 individually. The shorter wavelength population (495 nm excitation, 575 nm emission) remained stable up to 100 μM dopamine and was only quenched at higher concentrations when aggregation was observed (***Figure 3e***, green circles). By contrast, dopamine effectively quenched the fluorescence of the longer wavelength population (580 nm excitation, 650 nm emission) in the range of 5-50 μM dopamine (***Figure 3e***, red triangles). We conclude that the quench+shift phenomenon observed with UV excitation (***Figure 3d***) was due to the selective quenching of the longer wavelength population by the moderate amounts of dopamine used in our initial screen (50 μM, schematic in ***Figure 3a***). We therefore propose that the relative fluorescence of these two spectrally distinct S5-AgNC populations, under visible excitation, can be used as a fluorescent, ratiometric (*F*_575_/*F*_650_) and quantitative “turn-off” sensor (*F*_575_: invariant calibration, *F*_650_: indicator) of dopamine concentration *in vitro*.

### Specificity of silver nanoclusters for dopamine over known homologues and competitors

Encouragingly, both S4- and S5-templated silver nanoclusters were highly responsive to dopamine and relatively insensitive to the other neurotransmitters tested. However, we wondered if the sensors would be specific to dopamine over more closely related small molecules. We therefore interrogated both S4- and S5-stabilized AgNCs with catecholamine precursors and metabolites of dopamine (L-tyrosine, phenylalanine, L-DOPA, and DOPAC), as well as with small molecules known to interfere with dopamine sensing (L-tryptophan, and uric acid) (see ***Supplementary Figure S6*** for chemical structures).^10,26^

We tested whether these related molecules were capable of enhancing the formation of visible (615 nm emission) fluorescent nanoclusters by DNA sequence S4. Under 525 nm illumination, synthesis of S4-AgNCs in the presence of dopamine again produced a robust increase in 615 nm emission. However, synthesis in the presence of the other related molecules failed to significantly enhance fluorescent emission (***Figure 4a***). We further tested whether these other molecules were capable of quenching S5-AgNC fluorescence. Again, we observed that the addition of dopamine to existing S5-AgNC caused robust quenching of fluorescence (600 nm emission, 535 nm excitation) while S5-AgNC was unaffected by addition of other tested molecules (***Figure 4b***). We conclude that catecholamine precursors, dopamine metabolites, and known dopamine competitors do not significantly alter the fluorescence of these two DNA-AgNCs.

**Figure 4.**
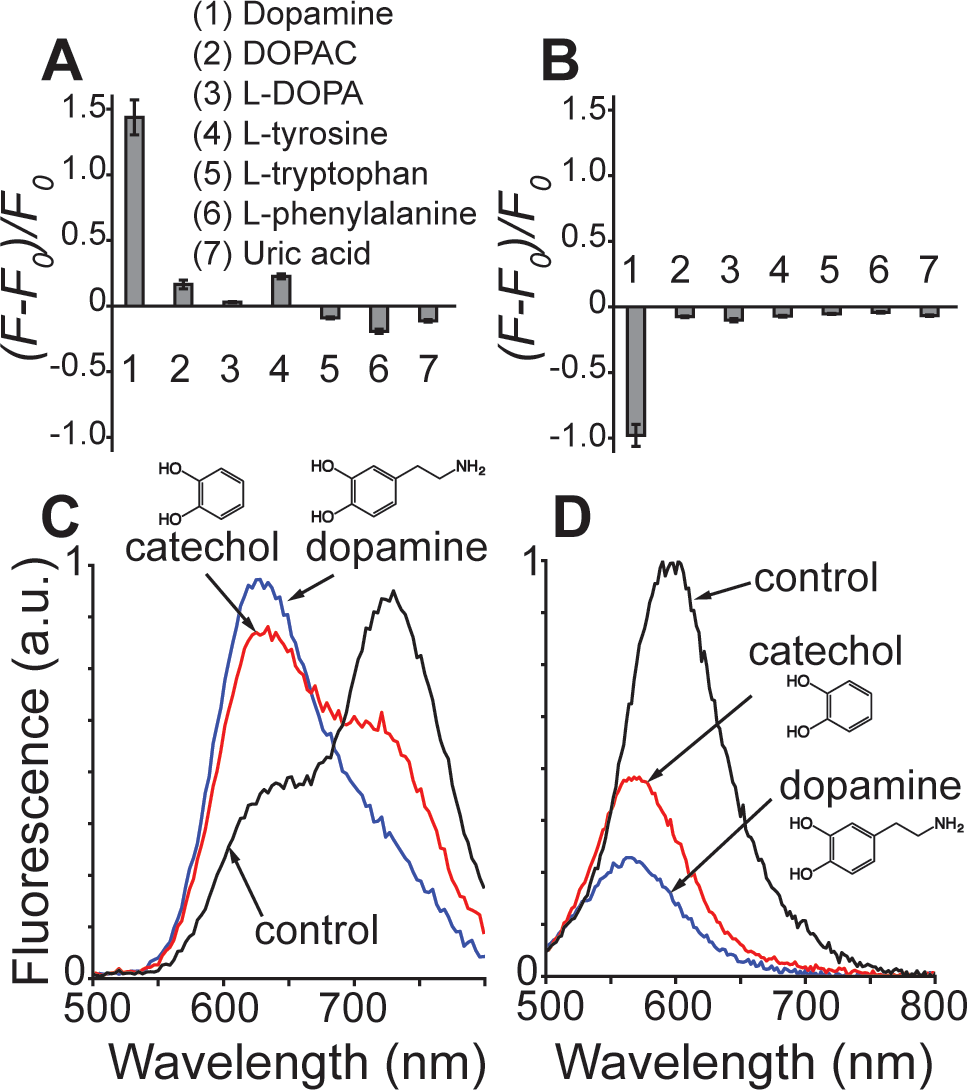
Preference of DNA-silver nanocluster sensors for dopamine over catecholamines and metabolites. **(a-b)**Change in fluorescence emission intensity,Δƒ = (*F-F_0_*)/*F_0_*, of nanoclusters assembled by templates **(a)** S4 or **(b)** S5 in the presence of dopamine, dopamine homologues or known competitors ([DNA] = 12.5 μM, [NT] = 50 μM). Both constructs showed high specificity for dopamine. **(c-d)** Fluorescence emission spectra of nanoclusters assembled by template S4 **(c)** or S5 **(d)** in the presence of dopamine, catechol or NT-free solution. Note: As in previous experiments, small molecules were added before synthesis of S4 (excitation: 525 nm, emission: 615 nm) **(a,c)** and after synthesis of S5 (excitation: 535 nm, emission: 600 nm) **(b,c)**.

To better understand the mechanism of dopamine interaction with DNA-silver nanoclusters, we investigated the role played by the catechol group in the interactions between dopamine and DNA-AgNCs. We compared the dopamine response of S5- and S4- templated AgNCs to the corresponding catechol response (***Figure 4c-d***). Again, we added dopamine, catechol, or control solution (buffer) prior to nanocluster synthesis for construct S4 and after nanocluster synthesis for construct S5. For both sensors, catechol modulated nanocluster fluorescence in a manner similar to dopamine, albeit to a lesser extent (***Figure 4c-d***). Like dopamine, catechol increased the formation of visible (615 nm emission) nanoclusters, while slightly decreasing the formation of nIR nanocluster (730 emission) (***Figure 4c***). Catechol also induced the quench+shift phenomenon, though to a lesser degree than dopamine (***Figure 4d***). However, dopamine had a more pronounced ability to prevent nIR nanocluster assembly (***Figure 4c***). This similarity suggests the catechol group is critical to the perturbation of DNA-AgNC fluorescence by dopamine. However, other molecules we tested containing a catechol group (namely DOPAC and L-DOPA) induced essentially no fluorescent change (***Figure 4a-b***).

We hypothesize that selectivity for dopamine is based on the small size of its alkylamine group. All other catecholamines tested contained a bulkier alkyl group (see ***Supplementary Figure S6*** for chemical structures), which could sterically hinder interactions with the DNA-AgNC complex. We presume that direct interactions between dopamine and the DNA strand perturb the DNA-silver nanocluster conformation.^16^ Dopamine binding to template S4 before nanocluster synthesis may destabilize the conformation necessary to form the near infrared population. Similarly, dopamine could destabilize the 650 nm nanocluster population formed by sequence S5 by perturbing the conformation of the DNA. However, dopamine could also promote direct conversion from one species to another with a different emission wavelength. Alternatively, dopamine may irreversibly interact with the DNA bases, resulting in cleavage or formation of adducts. In previous reports, quinones formed by the oxidation of dopamine were observed to quench the fluorescence of certain AgNCs, potentially through irreversible DNA damage.^27–30^ Such modifications to the DNA bases would produce changes in fluorescence by eliminating Ag+ binding domains or orbital interactions between DNA bases and the AgNC, or by perturbing a conformation critical to cluster formation or the continued stability of a cluster after synthesis.^12,28,29,31^ Elucidating the mechanism of dopamine action (and especially its reversibility) will be critical to establishing this technique as a viable option for *in vivo* monitoring.

We conclude that DNA-stabilized AgNCs can serve as selective, sensitive, ratiometric fluorescent sensors of dopamine concentration. Specifically, we identify stabilizing DNA sequences for fluorescent silver nanoclusters that support two complementary dopamine detection approaches *in vitro*: one (S4) provides a “turn-on” sensor that reports on the presence of dopamine during DNA S4-AgNC synthesis and the other (S5) provides a “turn-off” sensor that is quenched by the addition of dopamine to pre-formed DNA S5-AgNCs. Both DNA sequences produce two distinct emission peaks, thereby enabling quantification of dopamine concentration using ratiometric fluorescent sensing.

Ratiometric sensors offer important advantages over conventional fluorescent sensors, which report on the presence of a target molecule through a fractional change in overall emission intensity, Δƒ = (F-F_0_)/F_0_, at a single emission wavelength^32^. In conventional fluorescent sensing, the magnitude of the signal is often small and both baseline and target-modulated fluorescence are susceptible to interference (due to non-uniform concentration distribution, photobleaching, aggregation or nonspecific interactions with the environment, such as local pH changes,^33,34^) that complicates interpretation. Ratiometric fluorescent sensors, with their two spectrally distinct fluorescent signals, can be more responsive and less susceptible to interferences, especially if the fluorophores are chemically related and/or respond similarly to environmental conditions. DNA S5-AgNCs’ ratiometric signal is typical in that one of the emissions changes in response to target (dopamine) while the other remains invariant, and so can be used to scale or normalize the indicator signal. DNA S4-AgNCs’ ratiometric signal offers enhanced sensitivity because its two emissions have opposite dependency on dopamine.

With proper calibration, ratiometric sensing can report not merely the presence of and directional change in a target analyte, but also its concentration. Basal dopamine concentrations in relevant brain areas, including the extracellular matrix of the striatum, are commonly 1nM-1μM, though they can be higher in the vicinity of the synapse shortly after release.^7,35–38^ While the synthesis-dependent signal from DNA S4-AgNCs is sensitive in this range (10 nM - 100 μM), the sensitivity and working range of the post-synthesis responsive (and therefore *in vivo* compatible) DNA S5-AgNC nanosensor (1-100 μM) falls outside these commonly measured values. Ratiometric fluorescent sensors are generally challenging to engineer,^34^ but the sequence modularity of DNA-AgNCs offers a flexible new toolkit. Improvements in sensitivity might be achieved through directed mutation or automated screens of DNA sequence and nanocluster synthesis space. Given the documented successes of such screens^39^ as well as the ability to engineer FRET pairs,^40^ there is a real possibility of discovering clusters with sufficient sensitivities and temporal responses to enable real-time, quantitative imaging of neurotransmitters *in vivo*.

In summary, we present DNA-silver nanoclusters as novel, low-cost, easy-to-assemble, biocompatible reporters that offer selective, optical, quantitative sensing of dopamine concentration *in vitro*. With further development, DNA-stabilized AgNCs may enable spatially distributed, real-time, quantitative, optical monitoring of dopamine *in vivo*.

## METHODS

### Chemicals

Unless otherwise noted, chemicals were purchased from Sigma Aldrich (St. Louis, MO, USA): Dopamine hydrochloride, Acetylcholine chloride (≥99% TLC), Histamine dihydrochloride (≥99.0% AT), γ-Aminobutyric acid (GABA) (BioXtra, ≥99%), L-Aspartic Acid (reagent grade, ≥98% HPLC), Glycine (from non-animal source, meets EP, JP, USP testing specifications, suitable for cell culture, ≥98.5%), Epinephrine hydrochloride, L-Glutamic acid (*ReagentPlus*^®^, ≥99% HPLC), Uric acid (≥99%, crystalline), 3,4-Dihydroxy-L-phenylalanine (L-DOPA) (≥98% TLC), L-tryptophan (reagent grade, ≥98% HPLC)), L-Tyrosine (BioUltra, ≥99.0% NT), 3,4-Dihydroxyphenylacetic acid (DOPAC) (98%), L-Phenylalanine (reagent grade, ≥98%), (−)-Norepinephrine (≥98%, crystalline).

### Oligonucleotides

DNA oligos (detailed in ***Table 1***) were purchased from Integrated DNA Technologies, Inc. (Coralville, IA, USA) with standard desalting, hydrated with MilliQ deionized water (DI), and stored at −20°C in stock concentrations of between 100 μM and 1 mM. Sodium phosphate buffer was prepared by titrating solutions of sodium phosphate monobasic and sodium phosphate dibasic (Fisher Scientific, Waltham, MA, USA) to a pH of 7.3 and composite concentration of 200mM. DNA-containing solutions were prepared by combining 25 μL of 100 μM DNA solution with 50 μL of 20 mM sodium phosphate buffer, heating at 90°C for 5 min, then snap cooling by running the sample tube under cool DI water for 10 to 20 seconds.

### Cluster synthesis and neurotransmitter screening

Neurotransmitter and screening library chemicals were prepared as 10 - 100 mM aqueous stock solutions at 4°C. Silver nitrate (AgNO_3_, 99.99%), sodium borohydride (NaBH_4_, 99.99%, pellets) and sodium hydroxide (NaOH, >97.0%, pellets) solutions were prepared fresh for synthesis of silver nanoclusters (AgNC). Dopamine solutions were prepared fresh daily, and protected from light. ***Note:*** While some neurotransmitters were supplied as their HCl salt, we observed no systematic response of AgNC fluorescence to the presence of HCl during our initial screens, indicating that the responses we observed resulted from the neurotransmitter species and not interaction between Ag+ and Cl^−^. For **Method 1** (effect of NT on assembly of fluorescent DNA-AgNCs, **Figure 1b**), 50 μL of 100 μM solutions of each neurotransmitter (or DI) were added to DNA solutions (75 μL), and allowed to incubate for fifteen to twenty minutes at room temperature. To create nanoclusters, 12.5 μL of 0.16 mM silver nitrate were added to each sample, followed by 12.5 μL of freshly prepared 0.8 mM sodium borohydride diluted in 100 μM sodium hydroxide. Solutions were vortexed and left to incubate at 4°C for two hours. Then 20 μL of each DNA-AgNC/neurotransmitter solution was transferred to a 384-well plate (Product #3821BC, Corning, Inc.) for fluorescence measurements. The final concentration of DNA was 16.7 μM. For **Method 2** (effect of NT on assembled DNA-AgNC fluorescence, **Figure 1c**), 12.5 μL of 1.6 mM AgNO_3_ was added to the DNA solution and allowed to incubate in the dark for 10-20 minutes. Next, 12.5 μL of freshly prepared 0.8 mM NaBH_4_ diluted in 100 μM NaOH was added to the DNA-Ag solution. The solution was vortexed and left to react in the dark for 16-20 hours at 4°C. 10 μL samples of the resulting DNA-AgNCs were placed in a 384-well plate (Product #3821BC, Corning, Inc.) and combined with 10 μL of a 100 μM neurotransmitter solution or DI water as a control. The final concentration of DNA was 12.5 μM. Each sample was prepared in triplicate. The well plate incubated for 2-4 hours in the dark at 4°C until measurements were taken.

### Fluorescence detection

Fluorescence excitation and emission spectra of each sample were measured using a Tecan Infinite 200Pro plate reader with Tecan i-control software (Tecan Group Ltd.) between 1 and 5 hours after either chemical reduction or addition of NTs. Step size and bandwidth were set to 2 nm and 20 nm for emission scans, and to 5 nm and 5-10 nm for excitation scans, respectively. All scans were performed using Top Mode. The gain was kept constant at 105 (when using 260 nm excitation) or at 75 (when using visible excitation). Empty wells were processed to check for autofluorescence of the wellplate. For each DNA-AgNC, we recorded the proportional change in fluorescence, Δƒ = (F-F_0_)/F_0_, of a single emission wavelength in the presence of each NT (**Figure 1b-c**). We used the peak emission intensity at peak visible excitation in the absence of NTs (*F_0_*) and the emission intensity at the same excitation and emission wavelengths in the presence of NTs (*F*).

### SAFETY

No unexpected, new, and/or significant hazards or risks associated with the reported work were encountered.

## SUPPORTING INFORMATION

Supporting Information is available free of charge on the ACS Publications website. Fluorescence emission spectra of all DNA sequences (Supplementary Figure 1); distinct spectral populations for sequence S4 (Supplementary Figure 2) and sequence S5 (Supplementary Figure 4); ratiometric responses to dopamine concentration (Supplementary Figure 3); and structural formulas for neurotransmitters and small molecules in the screening library (Supplementary Figures 5 and 6) are provided.

## ABBREVIATIONS

DNA-AgNC: DNA-stabilized silver nanocluster
DOPAC: 3,4-Dihydroxyphenylacetic acid
L-DOPA: 3,4-Dihydroxy-L-phenylalanine
GABA: γ-Aminobutyric acid
NT: neurotransmitter

## Author contributions

J.T.D.O., D.K.F. and S.P. designed the research. J.T.D.O., A.T, J.W.H. and D.V. conducted experiments and performed data analysis. J.T.D.O., B.N.Q., D.K.F. and S.P. prepared and wrote the manuscript.

## ACKNOWLEDGMENT

The authors thank S. Helmy for helpful discussions.

